# Resting state EEG as a biomarker of Parkinson’s disease: Influence of measurement conditions

**DOI:** 10.1101/2020.05.08.084343

**Authors:** Henry Railo, Ilkka Suuronen, Valtteri Kaasinen, Mika Murtojärvi, Tapio Pahikkala, Antti Airola

**Affiliations:** Department of Clinical Neurophysiology, University of Turku, Turku, Finland; Turku Brain and Mind Centre, University of Turku, Turku, Finland; Department of Psychology, University of Turku, Turku, Finland; Department of Future Technologies, University of Turku, Turku, Finland; Clinical Neurosciences, University of Turku and Turku University Hospital, Turku, Finland

## Abstract

Resting state electroencephalographic (EEG) recording could provide cost-effective means to aid in the detection of neurological disorders such as Parkinson’s disease (PD). We examined how many electrodes are needed for classification of PD based on EEG, which electrode locations provide most value for classification, and whether data recorded eyes open or closed yield comparable results. We used a nested cross-validated classifier which included a budget-based search algorithm for selecting the optimal electrodes for classification. By iterating over variable budgets, we show that with eyes open recording, only 10 electrodes, localized over motor and occipital areas enable relatively accurate classification (AUC = .82) between PD patients (N=20) and age-matched healthy control participants (N=20). Classification accuracy only slightly increased when all 64 electrodes were included (AUC = .85). With the data recorded eyes closed, classification was not statistically significantly above chance even with full set of 64 electrodes (AUC = .55). These results show that classification based on small number of EEG electrodes is a promising tool for classifying PD, but measurement conditions and electrode locations can have a significant effect on classifier performance.

## 1. Introduction

Parkinson’s disease (PD) is a progressive degenerative neurological disorder whose cardinal behavioral signs are rest tremor, slowness of movement (bradykinesia), and muscular rigidity. The characteristic neuropathological feature of PD is the formation of abnormal alpha-synuclein aggregations, and the degeneration of dopaminergic cells in the midbrain region substantia nigra pars compacta. Although multiple biomarkers of PD have been proposed^29^, the diagnosis of PD is primarily based on motor symptoms. This is problematic because by the time the motor symptoms emerge, the neuropathological changes of PD are already widespread^10^, and a large proportion of dopaminergic cells has been lost^8^. An effective neuroprotection for PD will require earlier disease detection, and procedures that aid in the identification of PD from early biomedical signals are therefore needed. One of the proposed approaches is to classify PD based on electroencephalographic (EEG) recordings. EEG, which measures electrophysiological activity of large neuronal populations from the scalp, is low-cost, very easy to measure, and the recording systems are available in all hospitals. While studies have reported high accuracy in classifying PD from EEG, previous studies have not systematically examined how classification performance is affected by the number and location of EEG electrodes, and whether classification accuracy differs between eyes open or closed EEG recordings.

The degeneration of dopaminergic neurons in substantia nigra causes abnormal communication between neurons in the basal ganglia and cortex, and these changes are also visible in scalp recorded EEG^9,16,33^. PD “slows down” the EEG spectrum^4,21,26,28^. Studies which have classified PD patients and neurologically healthy age-matched controls based on linear features calculated from different frequency bands have yielded approximately 75–82% accuracy^5,6^. Building on the assumption that instead of reflecting linear stochastic processes, EEG is characterized by non-linear dynamics^18^, studies have also shown that non-linear methods provide valuable additional information when examining how EEG of PD patients and healthy controls differ^15,17,27^. In studies employing non-linear measures for classification, accuracy has been around 80–95%^13,15,17^. For instance, Lainscsek et al.^15^ showed that nine patients and nine age-matched controls could be classified almost perfectly with merely one second of 64-channel EEG when delay differential equations were used. Liu et al.^17^ studied 17 PD patients and 25 healthy controls using 10-channel resting eyes-closed EEG, and reported over 90% classification accuracy based on discrete wavelet transformation and sample entropy.

The aim of the present study was to examine three key aspects of classification of PD from EEG recordings. First, we wanted to examine how classification accuracy is influenced by the number of electrodes used. From the practical point-of-view it would be a considerable advantage if the classification of PD from EEG could be performed with minimal number of electrodes. This would make the EEG set-up fast, and high-density EEG montages are also rarely available in hospitals. Second, we wanted to examine if specific electrode locations provide most value for classification. If certain scalp locations provide more information for classification of PD than others, this information could be used to optimize EEG recording montages for PD classification. Our third aim was to examine if classification is influenced by whether the resting state EEG is recorded eyes open or closed. Because there are substantial differences in EEG dynamics between eyes closed and open conditions^2,12,13^—modulating, for instance, motor-related neural activity^7,25,32^—it could also affect classification performance. Most previous studies which classify PD based on resting state EEG have used eyes closed recordings^6,13,17^. Two studies which did not examine resting state (but visual event-related) EEG used eyes open recordings ^5,15^.

To examine how classification performance changes as a function of number of electrodes, we implemented a novel electrode-wise feature selection method which first fits a model to the features extracted from the single best electrode, and then augments the model further with features from additional performance-maximizing electrodes, until a pre-set budget for electrodes in the model is depleted. By iterating over variable budgets, we were able to observe at which number of electrodes the increase in classification performance becomes negligible, and which electrodes provide most value for classification. This procedure was run for eyes-closed and eyes-open resting state EEG data.

## 2. Materials and Methods

### 2.1. Participants

We tested 20 PD patients diagnosed using the UK Brain Bank criteria or MDS clinical criteria, and 20 age-matched control participants without neurological disease. One control participant was left-handed, others were right-handed. Demographical information, and clinical and neuropsychological measurements of the patients and participants are presented in Table 1. The severity of motor symptoms of PD was measured with the Movement Disorder Society Unified Parkinson’s Disease Rating Scale (MDS-UPDRS, Section III). Eight patients had MDS-UPDRS ≤ 20, which corresponds to average score at PD onset^14^. Seven patients had MDS-UPDRS 22–35, and the remaining five patients had MDS-UPDRS ≥ 35. The average rest tremor amplitude in the upper extremities was 1.1 (SD = 0.88; UPDRS item 3.17) in PD patients, indicating that patients’ tremor was < 1 cm in maximal amplitude. In five patients the tremor was left lateralized, and in four patients right lateralized (of the remaining patients six had no tremor, and five had bilateral tremor).

**Table 1.** Comparison of demographic, clinical and neuropsychological data between PD and control groups

|  |  |  |  |  |
| --- | --- | --- | --- | --- |
|  | **Control** | **PD** | **test statistic** | **p\*** |
| N | 20 | 20 | n/a | n/a |
| sex (male/female) | 8/12 | 9/11 | 0.10 | 0.74 |
| Age (mean +/- SD) | 68 (6.0) | 70 (7.2) | -0.90 | 0.37 |
| levodopa equivalent daily dose, mg (mean +/- SD) | n/a | 663 (509) | n/a | n/a |
| Disease duration, years (mean +/- SD) | n/a | 6 (4.9) | n/a | n/a |
| Mini-mental state score (mean +/- SD) | 28 (2.0) | 28 (1.8) | 0.62 | 0.53 |
| Beck's depression inventory | 5 (3.0) | 8 (6.2) | -2.07 | 0.04 |
| MDS-UPDRS, motor (mean +/- SD) | 5 (3.0) | 29 (16.4) | -6.02 | < .001 |
| * t statistic (continuous variables) or Chi-squared statistic (categorical data) |  |  |  |  |

When compared to the control participants, the patients had more depressive symptoms (as measured by Beck’s depression inventory, BDI), but otherwise the groups were comparable. Mini Mental-State (MMSE) questionnaire was used to assess cognitive impairment. The patients were asked to go through a voluntary medication break before the test session to counteract possible intervening effects of medication. Thirteen patients underwent the medication break, and did not take their morning medication during the day of the study (at least 12 h break in medication). These patients can therefore be considered being in OFF phase.

The study protocol was approved by the local Ethics Committee and followed the Declaration of Helsinki Ethical Principles for Medical Research Involving Human Subjects.

### 2.2. Electroencephalography: recording and preprocessing

EEG was measured with 64 active electrodes with sampling rate of 500 Hz, and 0.16–125 Hz filtering during recording using a NeurOne Tesla amplifier. In addition, eye movements were measured from electrodes placed beside and below the right eye. Electrode impedances were brought close to 5kΩ at the beginning of the measurement. Two minutes of eyes-open and two minutes of eyes-closed resting state EEG was collected per participant (except, due to a technical failure, the first patient’s eyes-closed data was lost). During the data collection session, the participants and patients also took part in an experiment investigating speech deficits in PD^23^.

Data preprocessing was performed using a standardized and fully automated PREP pipeline^3^. During preprocessing, 50 Hz line noise was removed, bad channels were interpolated, and EEG was referenced to robust average reference. The datasets are available for download at https://osf.io/pehj9/.

### 2.3. Feature extraction

Classification was based on 64 EEG electrodes (electrodes used to measure eye movements were not included). We applied a third order Butterworth band-pass filter to separate four frequency bands corresponding with delta, theta, alpha and beta rhythms. The respective lower and upper limits of the aforementioned frequency bands were [0.5, 4.0], [4.0, 8.0], [8.0, 13.0] and [13.0, 30.0], the unit of measure being Hz.

We computed the sample entropy (SampEn^24^) for each such frequency band resulting in 256 numeric features per subject. We decided to use SampEn because it^17^ and other similar measures that are sensitive to non-linear dynamics of EEG ^13,15^ have been previously reported to work well with PD classification. SampEn is a measure used for quantifying the unpredictability of a physiological time series, based on estimating the probability that two matching sequences in the time series of a given length *m* are still similar at length *m*+1. It is closely related to another complexity measure known as approximate entropy (ApEn), but designed to eliminate the bias that originates from ApEn counting each sequence in the time series as matching itself. Thus, SampEn gives a less biased measure of similarity. Contrary to ApEn, SampEn is also independent on the sample length.

SampEn is defined by a series of *N* data points *X = x*_*1*_, *x*_*2*_, *…, x*_*N*_, an embedding dimension *m* and a tolerance *r*. The embedding dimension, also known as order, is used for constructing a set of template vectors *u*_*m*_(*i*) *=* [*x*_*i*_, *x*_*i*+*1*_, *…x*_*i*+*m*−*1*_] for each i msuch that 1 ≤ *i* ≤ *N* − *m* in the embedding space **R**^m^. Another such set *u*_*m*+*1*_(*i*) of template vectors is constructed in the embedding space **R** ^m+1^. Also, a distance function *d*(*u*(*i*), *u*(*j*)) is defined for template vectors *u*(*i*), *u*(*j*) as the Chebyshev distance (or, less often, some other distance metric) in their respective embedding spaces. Given a time series *x*(*n*), a distance metric *d*, an embedding dimension *m* and a tolerance *r*, SampEn is defined as

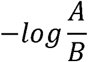

where

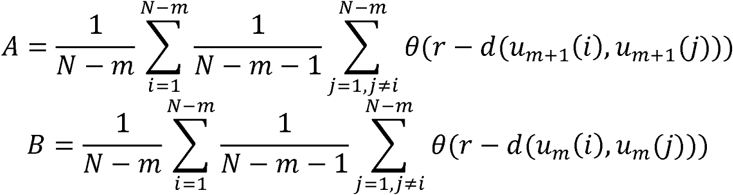

and where θ is the Heaviside step function.

In our experiment, the sample entropy was computed from a sample of 20 000 data points (40 seconds of EEG) using an embedding dimension of 2 and a filtering level of 0.2 * *σ*, where *σ* is the standard deviation of the sample.

### 2.4. Classification

For classification we used a logistic regression classifier (with a small constant l2-penalty to speed up the convergence) and estimated the performance of the classifier using a nested stratified 10-fold cross-validation algorithm. The rationale for using nested cross-validation instead of regular non-nested cross-validation when performing model selection is that performing model selection using non-nested cross-validation has been reported to result in highly biased performance estimate. Using nested cross-validation for model selection and performance estimation reduces this positive bias, yielding a more realistic estimate of true classification performance^30^. We estimated the classification performance for locally optimal subsets of features extracted from variable numbers of electrode channels, ranging from one electrode to a complete set of 64 electrodes. For finding such locally optimal subsets of electrodes for each subset cardinality, we used a custom budget-based greedy forward search algorithm^20^. We repeated the nested cross-validation procedure ten times in total, each time with a different random state used for dividing the data into training, validation and testing folds, and then averaged over the test performance estimates in order to mitigate the effect of chance on the performance estimates.

### 2.5. Budget-based electrode-wise forward search

To find the locally optimal sets of electrodes for training the model with given variable budgets we used a custom budget-based search algorithm. Given a budget B, the algorithm will iteratively select in total B electrodes, so that the validation performance estimate will be maximized for each new electrode chosen. New electrodes will be selected in a descending order with respect to the validation performance estimate, so that the electrode that the algorithm will select in each iteration is the one the introduction of which into the model will have the highest positive effect (or the least negative effect) on the validation performance estimate. The selected electrode will then be removed from the set of available electrodes, the cost of the model will be increased by one, and the selection procedure will continue until the budget is depleted. When the budget is depleted, the model will be trained using the selected set of electrodes and its performance will be estimated using the test fold. For an estimate of performance over the whole dataset, we used an averaging procedure. We used the estimate for area under the receiver operating curve (AUC) as the primary criterion when selecting new electrodes.

The analysis was performed in Python (script is available at https://gitlab.utu.fi/ilksuu/pd_package) with Numpy^31^, Scikit-learn^22^ and EntroPy^19^ as external packages.

### 2.6. Statistical significance of classification performance

To determine if classification performance was higher than predicted by chance, we performed permutation tests with 10,000 permutation on the data with full set of electrodes, using leave-pair-out cross-validation (LPO) for minimal negative bias. Permutation test is an often-used method for assessing the reliability of a classifier by calculating a measure of probability that a classification performance equal or higher than that of the classifier under scrutiny can be achieved by chance ^11^. LPO is a special case of k-fold cross-validation for binary classification, in which every possible pair of instances from different classes are used as test folds. LPO provides nearly unbiased performance estimates, and is suggested as the preferred cross-validation scheme for estimating conditional AUC^1^. As a downside, LPO has a relatively high computational cost, which is why we chose to substitute it with ten iterations of randomized 10-fold cross-validation for the actual analysis.

## 3. Results

The comparison of EEG frequency spectrums in the eyes closed and eyes open condition is shown in Figure 1A and 1B. As expected, the eyes closed condition (orange line) showed stronger alpha (8–10 Hz) activity when compared to the eyes open condition (blue) in both control participants and PD patients, although this effect is weaker in PD patients. Figures 1C and 1D show the comparison of patient and control groups in eyes closed and open conditions, respectively using independent samples t tests. As previously reported^4,21,26,28^, EEG power was higher in the PD group in low frequencies (positive t values), and lower in high frequencies (negative t values), when compared to the control participants. The eyes closed and open conditions showed relatively similar effects, except that the increase of low frequency EEG power in PD patients around 2–4 Hz is stronger in the eyes closed condition than in eyes open condition. The decrease in high frequency EEG power in PD group is somewhat right lateralized (see Fig. 1C and 1D topography of t values).

**Figure 1.**
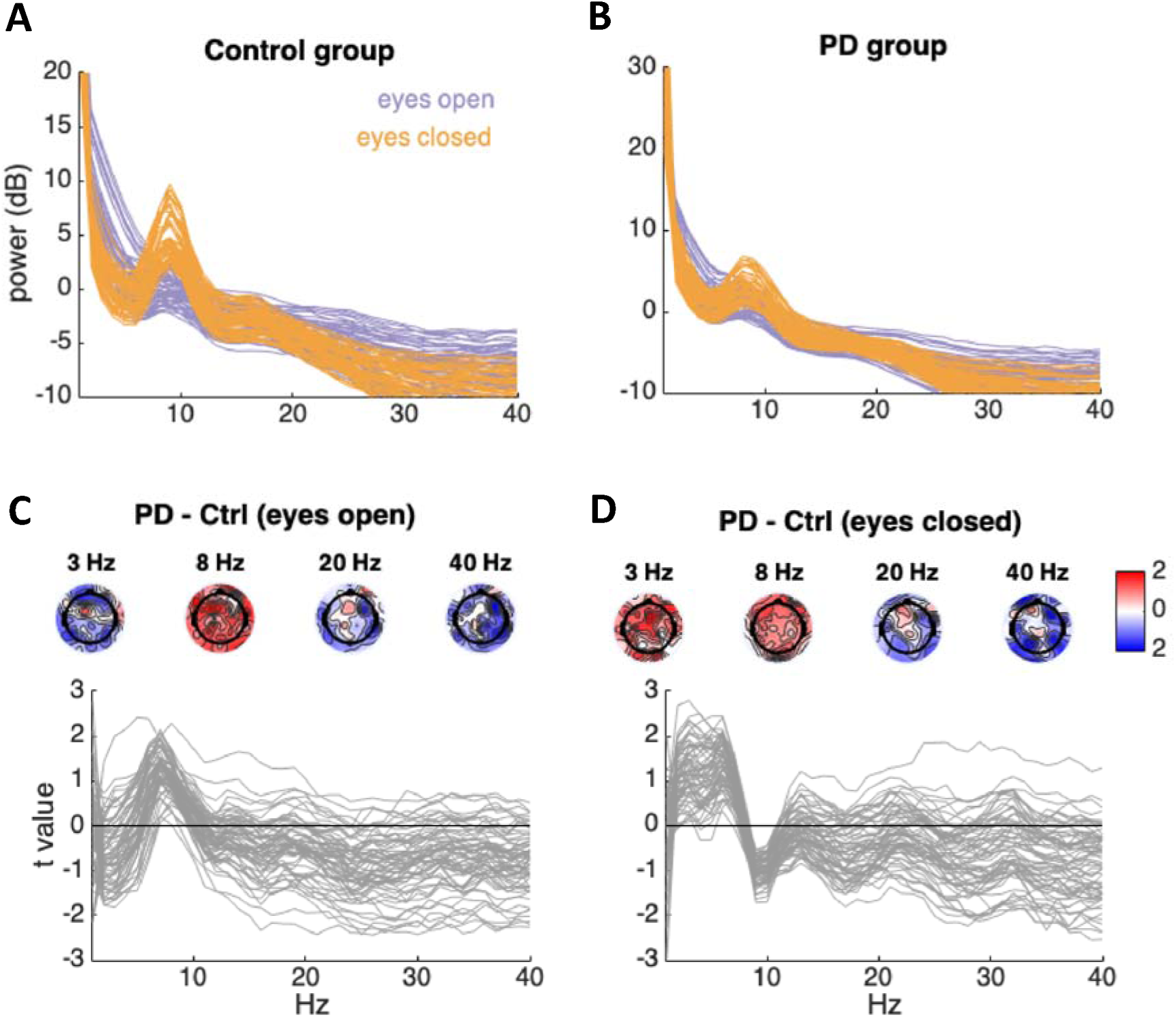
Overview of measured EEG signal. A) Power spectrum in eyes closed (orange) and open (blue) condition in the control group. Each line in the plot represents one channel. B) Power spectrum in eyes closed (orange) and open (blue) condition in the PD group. C) Comparison of EEG power between the patient and control groups in the eyes closed condition. The lines show the result (t statistic) of independent samples t tests, performed for each electrode. The scalp maps show the scalp distributions of the t values at four specified frequencies. D) Comparison of EEG power between the patient and control groups in the eyes open condition. In both panels C and D negative t values indicate that EEG power is lower in the PD group when compared to the control participants. The scalp maps show the distribution of t values. Color scale ranges from t = −2 (blue) to t = 2 (red).

Classification performance as a function of budget in the eyes open and closed conditions is shown in Figure 2. The orange lines represent AUC and the blue line is the proportion of individuals who were correctly categorized (accuracy, ACC). In the eyes open condition classification performance increases as the function of electrodes until around 10 electrodes (AUC = 0.823, ACC = 0.778, sensitivity = .83, specificity = .72). After this point additional electrodes only slightly improve the classifier. The permutation test demonstrated statistically significant classification in the eyes open condition (with 64 electrode budget) with AUC estimate of 0.855 (p = .0002; sensitivity = .85, specificity = .79). The permutation test showed that classification was not statistically significantly higher than chance-level the eyes closed condition (with 64 electrodes, AUC = 0.56, p = .36; sensitivity = .39, specificity = .68).

**Figure 2.**
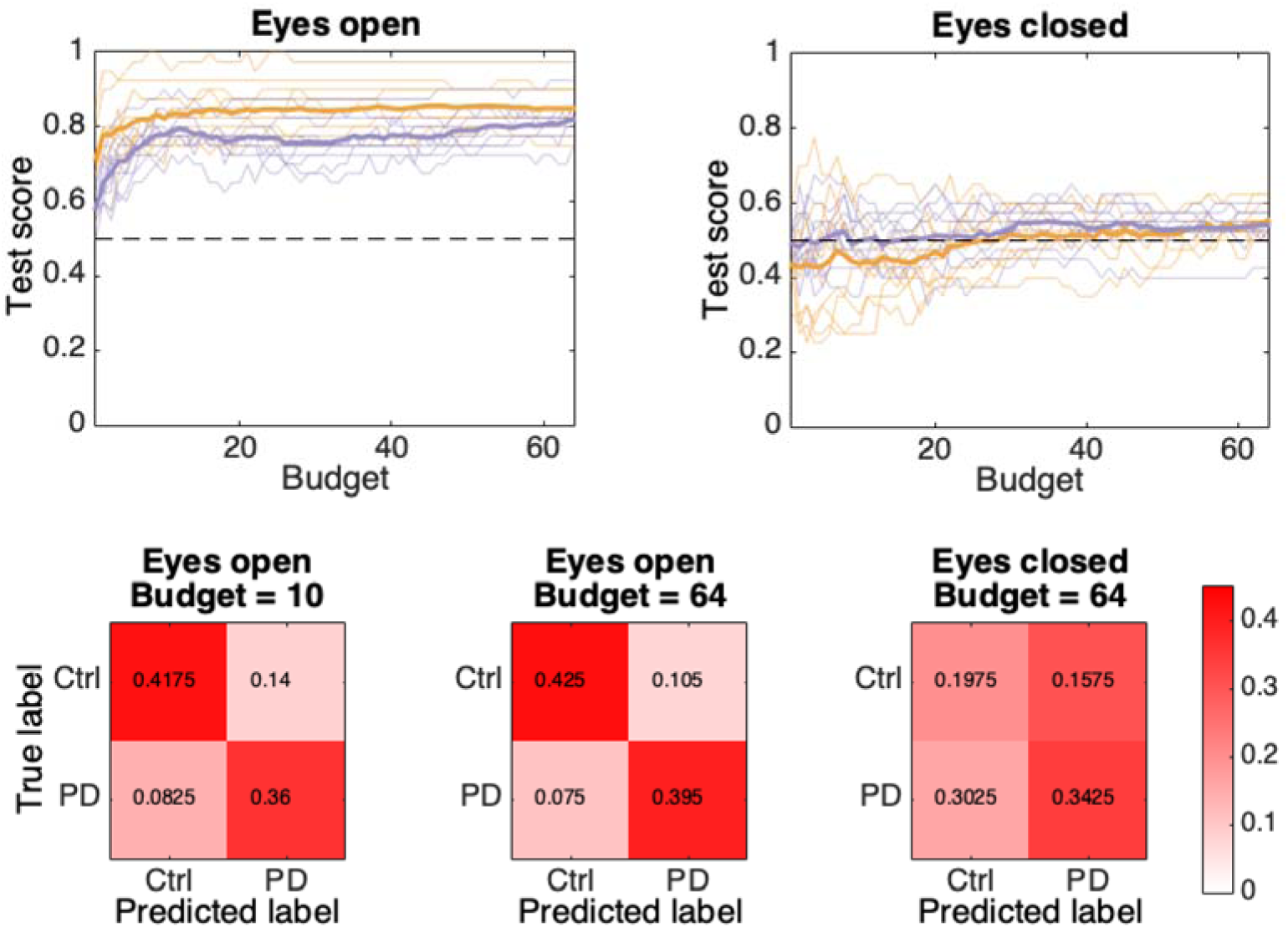
Classification performance as a function of number of electrodes used for fitting the model (i.e. budget) in the eyes open and closed conditions. The orange lines show the AUC and the blue lines the accuracy of correctly categorizing an individual. Thinner lines represent different training, validation and testing folds (thicker lines are their average). Confusion matrices from classifiers with 10 electrode budget with eyes open data, full 64 electrode budget with eyes open data, and full 64 electrode budget with eyes closed data are shown in the bottom panels. The confusion matrices have been calculated based on all 10 different random folds.

Certain electrodes were more likely to be selected for the nested cross-validation than others in the eyes open condition. Observing the pooled frequencies for a given electrode to be selected (eyes open condition), given a budget of 5, 10, 20, and 30 electrodes, is shown in Figure 3. When budget was 5, it was much more likely that left hemisphere electrode locations were selected into the model. With higher budgets, electrodes located over the motor cortex and occipital pole were most likely to be selected into the model, suggesting that these areas contained valuable information for the classifier. Prefrontal and parietal electrode locations were least likely to be selected into the model. Similar data is not presented for the eyes closed condition because classification was not successful even with full 64 electrodes.

**Figure 3.**
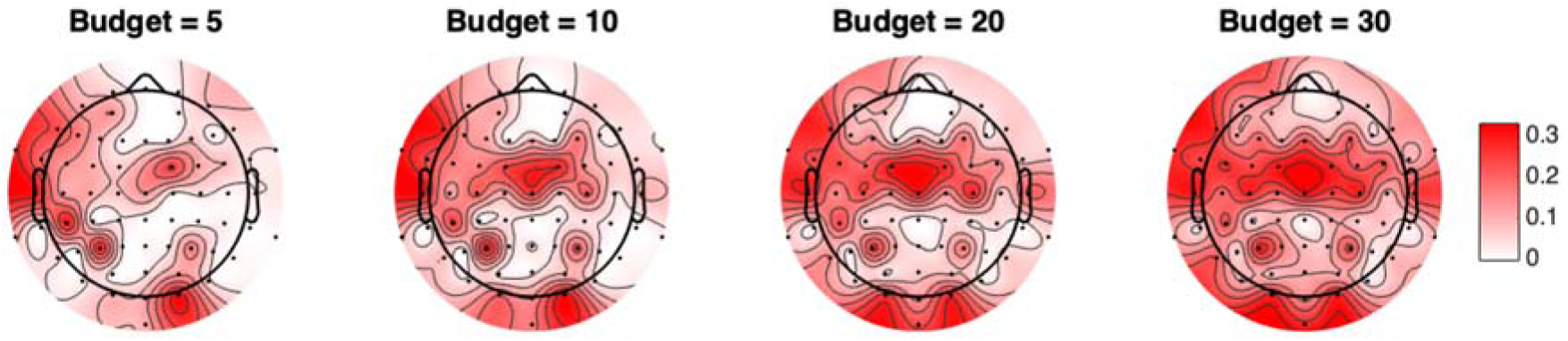
Probabilities that a given electrode was selected into the classifier in the eyes open condition given different budgets.

## 4. Discussion

Using a novel budget-based greedy search algorithm, we examined how well PD can be classified with variable numbers of electrodes based on resting state EEG data. Our results show that early/intermediate stage PD could be classified with relatively high sensitivity from resting state EEG data recorded eyes open with only roughly 10 electrodes. Higher number of electrodes did not significantly improve the classifier. The electrode locations that were most beneficial for the classifier were located over motor and occipital cortex. Interestingly, classification was not possible with eyes closed resting state data even when all 64 electrodes were included in the model. Altogether our results show that even low-density EEG is a promising tool for PD detection, but that recording conditions can significantly influence classification performance.

The failure to classify PD based on eyes closed data in the present study should not be taken to imply that PD cannot be classified by relying on eyes closed data. Failure to classify PD based on eyes closed EEG may indicate that the feature extraction method or the classification algorithm used in the present study was ill-suited for eyes closed data. Previously, Liu et al.^17^ showed that PD could be accurately predicted from eyes closed resting state EEG data when sample entropy was used for feature extraction. While this is at odds with the present results, direct comparison of the studies is difficult. Liu et al.^17^ do not report how severe PD the patients in their sample had. If the sample consisted of PD patients with e.g. severe rest tremor, high classification accuracy is relatively trivial as movement strongly affects EEG.

We chose to include only early or intermediate PD patients with relatively low levels of rest tremor to make the test clinically more realistic. If classifiers are to help in the diagnosis in future, naturally the classifiers should work with early stage PD patients. However, because our patients were all on medication, we cannot rule out confounds related to medication. Gomez et al. ^13^ showed that early non-medicated PD patients could be classified from healthy controls with high accuracy based on eyes closed resting state magnetoencephalography (MEG) with Lempel-Ziv complexity (however, the authors do not report patient characteristics such as UPDRS scores). The difference to present results could be related to different means of recording neural activity, and different feature extraction method.

Our data shows that electrodes over the motor cortex were valuable for the classifier, suggesting that activity in the motor cortex was a key contributor for the classification. Motor cortex activity is modulated by whether eyes are open or closed ^7,25,32^, and PD, of course, is characterized by motor symptom. Electrodes located in the occipital pole also provided valuable information for the classifier, suggesting that visual cortical activity can be used to differentiate PD from healthy controls. Note that these conclusions about possible *cortical* generators are indirect: EEG signal recorded from scalp does not afford accurate localization of neural activity because of volume conduction. Yet, our results do show that optimal classification may be afforded by EEG recording montages where electrodes are placed especially over motor and occipital cortices.

Previous studies have also shown that relatively high classification performance can be reached with ten electrodes^17^ and short recordings^15^, but the present study is the first that directly examined how the number of electrodes influences classification. Our results should not be taken to mean that additional recording channels cannot improve classification—for example, coupled with source reconstruction, high density EEG montages could provide benefit—but our results show that high-density EEG does not directly imply improved classification relative to low-density electrode montages. The locations where the electrodes are placed may be more important than number of electrodes. However, strong conclusions should not be drawn solely based on the present, relatively small sample. While our study is the first to use a budget-based classification algorithm, there is relatively large variation between previous studies^13,15^ in which brain areas allowed highest classification accuracy.

To conclude, given that also many previous studies have demonstrated high classification accuracy of PD based on EEG data^5,6,15,17^, the evidence suggests that EEG can be a valuable source for PD classification. The present results draw attention to the fact that measurement conditions can significantly affect classification performance.

## 5. Acknowledgments

The research was funded by Academy of Finland (H.R, grant #308533, and T.P. grants #313266 and #311273).

